# Intranasal HD-Ad Vaccine Protects the Upper and Lower Respiratory Tracts of hACE2 Mice against SARS-CoV-2

**DOI:** 10.1101/2021.04.08.439006

**Authors:** Huibi Cao, Juntao Mai, Zhichang Zhou, Zhijie Li, Rongqi Duan, Jacqueline Watt, Ziyan Chen, Ranmal Avinash Bandara, Ming Li, Sang Kyun Ahn, Betty Boon, Natasha Christie, Scott Gray-Owen, Rob Kozak, Samira Mubareka, James M. Rini, Jim Hu, Jun Liu

## Abstract

The COVID-19 pandemic has affected more than 120 million people and resulted in over 2.8 million deaths worldwide. Several COVID-19 vaccines have been approved for emergency use in humans and are being used in many countries. However, all of the approved vaccines are administered by intramuscular injection and this may not prevent upper airway infection or viral transmission. Here, we describe intranasal immunization of a COVID-19 vaccine delivered by a novel platform, the helper-dependent adenoviral (HD-Ad) vector. Since HD-Ad vectors are devoid of adenoviral coding sequences, they have a superior safety profile and a large cloning capacity for transgenes. The vaccine (HD-Ad_RBD) codes for the receptor binding domain (RBD) of the SARS-CoV-2 spike protein and intranasal immunization induced robust mucosal and systemic immunity. Moreover, intranasal immunization of K18-hACE2 mice with HD-Ad_RBD using a prime-boost regimen, resulted in complete protection of the upper respiratory tract against SARS-CoV-2 infection. As such, intranasal immunization based on the HD-Ad vector promises to provide a powerful platform for constructing highly effective vaccines targeting SARS-CoV-2 and its emerging variants.

## Introduction

The COVID-19 pandemic, caused by severe acute respiratory syndrome coronavirus 2 (SARS-Cov-2)^1^, is ongoing and has resulted in more than 120 million confirmed cases and at least 2.8 million deaths worldwide. The development of safe and effective vaccines against SARS-CoV-2 is a global health priority and a number of vaccine platforms have already been tested^2^. These include inactivated virus^3,4^, naked DNA delivered by electroporation^5^, mRNA delivered by lipid nanoparticles^6–9^, viral vectors such as nonreplicating adenovirus^10–17^, replication-competent vesicular stomatitis virus (VSV)^18^ or yellow fever virus^19^, and recombinant protein delivered by nanoparticles^20,21^. To date, at least four vaccines have been evaluated in phase III clinical trials and results have been published, including the mRNA vaccines from Pfizer-BioNTech^22^ and Moderna^23^, the ChAd vaccine from AstraZeneca^24^, the rAd26 and rAd5 vaccines from Russia^25^. The two mRNA vaccines and the Janssen rAd26 vaccine have received Emergency Use Approval by the FDA.

With the roll out of approved vaccines in many countries, several limitations of the first-generation of COVID-19 vaccines became increasingly apparent. The requirement of ultra-cold temperature for transportation and storage of the mRNA vaccines is a major limitation especially for global distribution in developing countries. In addition, all the approved vaccines are delivered by intramuscular injections, which may not be able to prevent the infection at the upper respiratory tract and stop viral shredding and transmission^26^.

Here, we investigate the performance of a COVID-19 vaccine developed in a novel platform, the helper-dependent adenoviral (HD-Ad) vector. HD-Ad, also known as gutless adenovirus, is the latest (third) generation of adenoviral vectors and has primarily been used for *in vivo* gene delivery in preclinical tests for the treatment of inherited genetic diseases^27,28^. HD-Ad was constructed with all the adenoviral coding sequences deleted and this eliminates the expression of unwanted adenoviral proteins^27,29^. This characteristic minimizes the host immune response to the vector and allows for long-term expression of the transgene in host tissues or organs^30–34^. In addition, HD-Ad does not integrate into the host genome, thereby eliminating the risk of introducing chromosomal mutations. Besides its excellent safety profile, HD-Ad has a high cloning capacity (up to 36 kb) for transgenes, which makes it possible to deliver large genes or multiple genes. All these features make HD-Ad an attractive platform for vaccine construction and delivery. In this study, we delivered the receptor binding domain (RBD) of the SARS-CoV-2 spike protein (S protein) to mice intranasally using an HD-Ad vector and examined its immunogenicity and protective efficacy.

## Results

### Cloning and expression of a secreted form of the SARS-CoV-2 RBD

The RBD sequence of SARS-CoV-2 was codon optimized (Table S1) and expressed from the chicken beta-actin (CBA) promoter with a cytomegalovirus (CMV) enhancer, to increase the transcription, and the first intron of the human *UbC* gene to increase mRNA stability (**Fig. 1A**). A DNA sequence encoding the 20-amino acid signal peptide of the human cystatin S protein was included upstream of the coding sequence of the RBD, which allows for RBD secretion. The BGH poly A tail was used to terminate the transcription.

**Figure 1.**
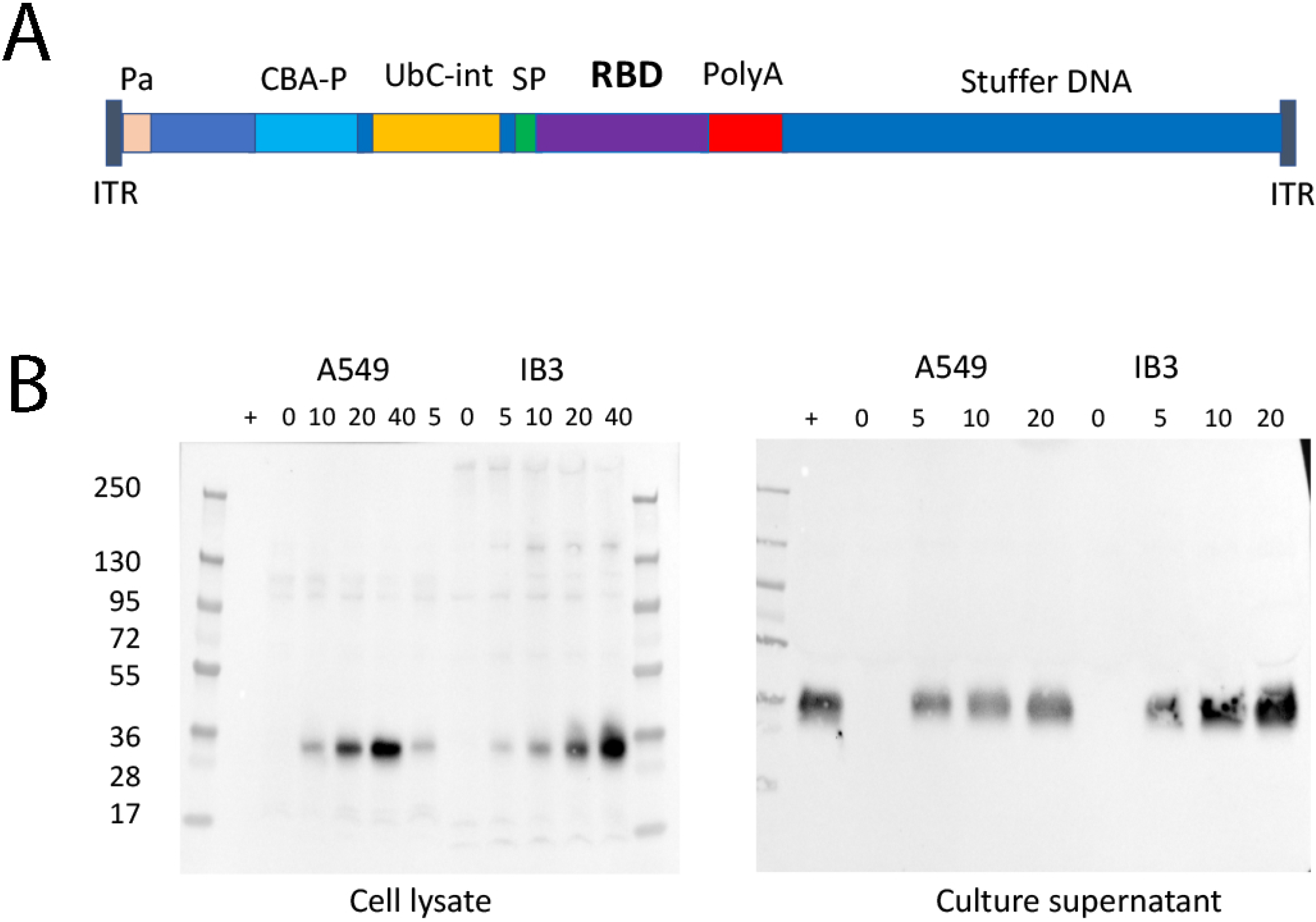
Construction of HD-Ad_RBD vaccine. **(A) Scheme of the HD-Ad_RBD vaccine.** ITR, inverted terminal repeat of the adenovirus; Pa, packaging signal of adenovirus; CBA-P: Chicken beta actin gene promoter with CVM enhancer; UbC-int, the first intron of human ubiquitin C gene; SP, the human cystatin-S signal peptide required for protein secretion; RBD, receptor binding domain of the spike protein; PolyA: transcription termination signal of the Bovine Growth Hormone gene; Stuffer DNA, noncoding human DNA used to make the vector genome size big enough for packaging. (**B**) **Western blot analysis of RBD expression and secretion**. Epithelial cells A549 and IB3 were transfected with HD-Ad_RBD at indicated dosages and cell lysates and culture supernatant were prepared and subjected to Western blot analysis using anti-RBD antibodies.

To examine the expression and secretion of the RBD, epithelial cells (A549 and IB3) were transfected with HD-Ad_RBD at different dosages and cell lysates and culture supernatants were prepared and subjected to Western blot analysis. The result showed that the RBD was expressed at high levels in both cell lines and in a dose-dependent manner (**Fig. 1B**). Importantly, we estimated that about 90% of the RBD was detected in the culture supernatant, indicating that the majority of the RBD was secreted into the culture media. The secreted transgene product delivered by HD-Ad vectors is known to reach both airway fluid and the blood circulation system^28^, thus the secreted RBD is expected to reach antigen presenting cells, locally and systemically, to induce antigen-specific immune responses.

### HD-Ad_RBD induces robust mucosal and systemic immunity

To examine the immune responses induced by HD-Ad_RBD, we intranasally immunized BALB/c mice (*n*=5) with three different doses of HD-Ad_RBD; three weeks later these mice were sacrificed and sera collected (**Fig. 2A**). ELISA analysis of the sera showed high levels of RBD-specific IgG in all three groups of mice immunized with HD-Ad_RBD, whereas low, if any, levels of IgG were detected in mice immunized with the HD-Ad control, which does not carry a transgene for protein expression (**Fig. 2B**). Animals vaccinated with 10^8^, 5×10^9^ and 10^10^ HD-Ad_RBD viral particles had reciprocal geometric mean titers (GMT) of 30,314, 459,479, and 378,929, respectively. This result indicates a dose-dependent response and that 5×10^9^ of HD-AD_RBD is the optimal dosage for vaccination.

**Figure 2.**
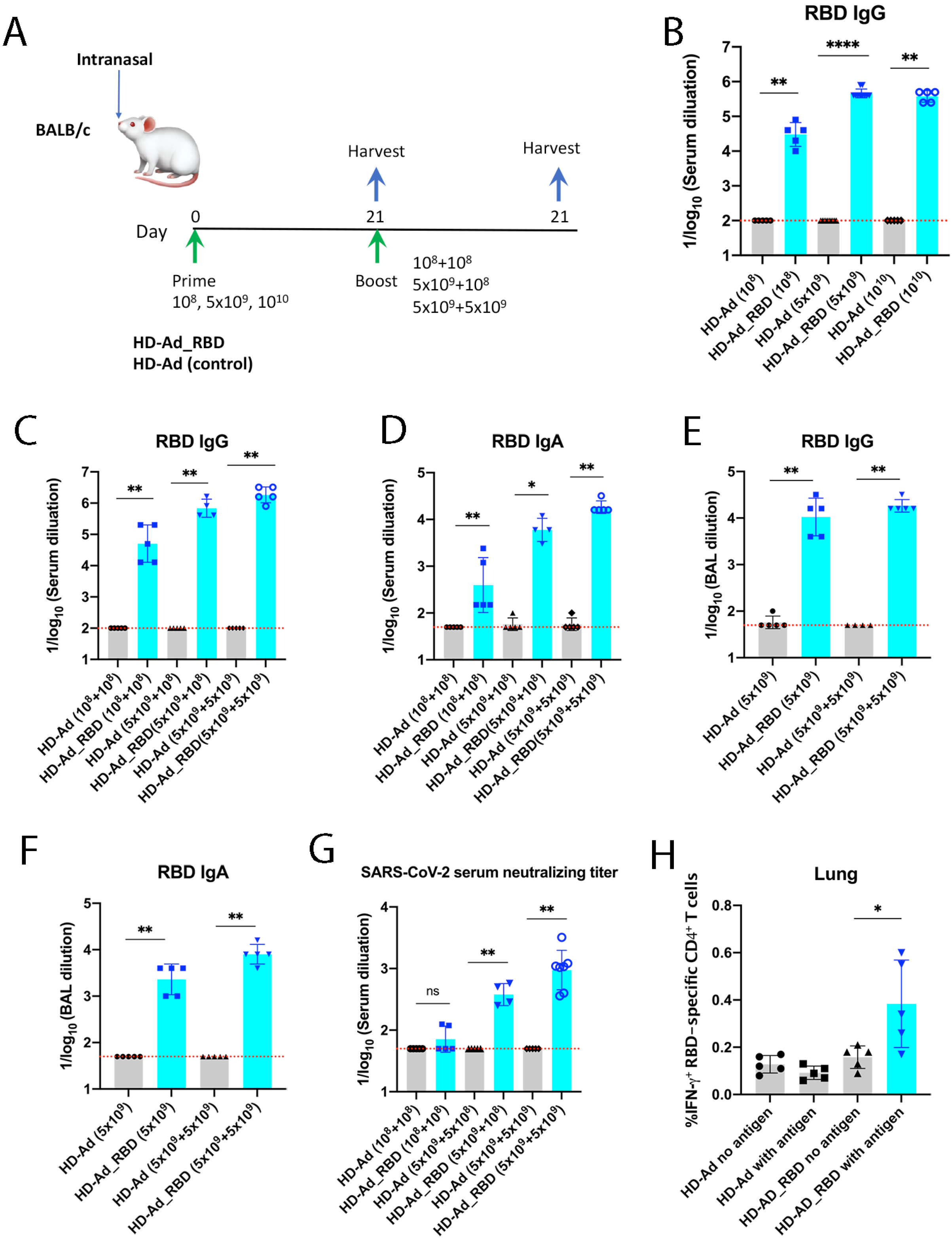
HD-Ad_RBD induces high levels of IgG, IgA and neutralizing antibody. (**A**) **Scheme of the experiments**. BALB/c mice were immunized with HD-Ad_RBD or HD-Ad via an intranasal route. (**B**) **Antibody responses of single vaccination with different doses**. RBD-specific IgG in sera were measured by ELISA. (**C & D**) Antibody responses of prime-boost vaccination with different doses. RBD-specific IgG (**C**) and IgA (**D**) in sera were measured by ELISA. (**E & F**) **Antibody responses detected in BALs**. BALs from single (5×10^9^) or prime-boost vaccinated (5×10^9^+5×10^9^) mice were collected and RBD-specific IgG (**E**) and IgA (**F**) were measured by ELISA. (**G**) Neutralizing activity of sera against SARS-CoV-2 were measured. (**H**) **Intracellular cytokine (IFN-γ) staining and flow cytometry analysis of CD4^+^ T cells**. Harvested lung cells were stimulated with or without purified RBD (10 μg/ml) for 12 h at 37°C and subjected to ICS analysis. In all figures, each dot represents an animal. Bars and errors represent the geometric mean with geometric SD. The red dotted lines indicate the limit of detection (LOD) of assays. Statistical analyses were performed by Mann-Whitney test: *p < 0.05; **p <0.01; ***p < 0.001.

We next tested the prime-boost intranasal vaccination regimen. Based on the results of the single dose vaccinations, we chose 10^8^ and 5×10^9^ HD-Ad_RBD viral particles for the prime immunizations, followed 3 weeks later by boost vaccinations at the same dosage (**Fig. 2A**). A lower boost dosage (10^8^ viral particles) was also tested in the 5×10^9^-primed group. Three weeks after the second vaccination, the animals were sacrificed and sera and bronchoalveolar lavages (BALs) were collected.

Boosting with 10^8^ HD-Ad_RBD viral particles increased the IgG titer ~1.5-fold relative to the single 10^8^ vaccination, and the IgG reciprocal GMTs for the 10^8^ prime-boost group, and the 5×10^9^-prime/10^8^-boost group were 50,476 and 688,862, respectively (**Fig. 2C**). Remarkably, the IgG reciprocal GMT in the 5×10^9^ prime-boost group increased 4-fold compared to that of the single vaccination, reaching 1,837,920 (**Fig. 2C**).

High levels of RBD specific IgA were also detected in the sea of the boosted animals and the highest level was detected in the 5×10^9^ prime-boost group, reaching a reciprocal GMT of 18,379 (**Fig. 2D**).

We also detected high levels the RBD-specific IgG and IgA in BLAs. For the 5×10^9^ prime-boost group, the IgG and IgA reciprocal GMTs were 18,379 and 8,000, respectively (**Fig. 2E & F**).

Similarly, there was a dose-dependent increase in the neutralizing activity of the sera against the SARS-CoV-2 virus (**Fig. 2G**). The reciprocal 50% inhibition dilution (ID_50_) GMTs of the neutralizing antibody in the 5×10^9^-prime/10^8^-boost group and the 5×10^9^ prime-boost group were 378 and 948, respectively.

Finally, we also detected an increase of IFN-γ producing CD4^+^ T cells in the lungs of animals vaccinated with 5×10^9^ HD-Ad_RBD viral particles (prime and boost) compared to the control groups, indicating that the Th1 response was activated (**Fig 2H**).

### HD-Ad_RBD protects hACE2 mice against SARS-CoV-2 infection

To examine the protective efficacy of HD-Ad_RBD, we intranasally immunized K18-hACE2 transgenic mice with HD-Ad_RBD (*n*=17) or the HD-Ad vector alone (sham control, *n*=18). SARS-CoV-2 does not infect regular mice due to the lack of the right receptor. The K18-hACE2 transgenic mouse model was generated by McCray *et al*.^35^ by expressing the human ACE2, the receptor for SARS-CoV-2, using the K18 gene expression cassette developed by one of our laboatories^36^. The hACE2 mice received a prime immunization of 5×10^9^ viral particles of HD-Ad_RBD or HD-Ad at day 1. At day 21, the mice received a boost immunization of the same dose of the vaccine or the control. At day 21 after the second immunization, hACE2 mice were intranasally challenged with 10^5^ 50% tissue culture infectious dosage (TCID_50_) of SARS-CoV-2. At day 1, 3 and 5 post-infection, mice were euthanized and lungs, spleen and heart were harvested for viral burden and cytokine analysis (**Fig. 3A**).

**Figure 3.**
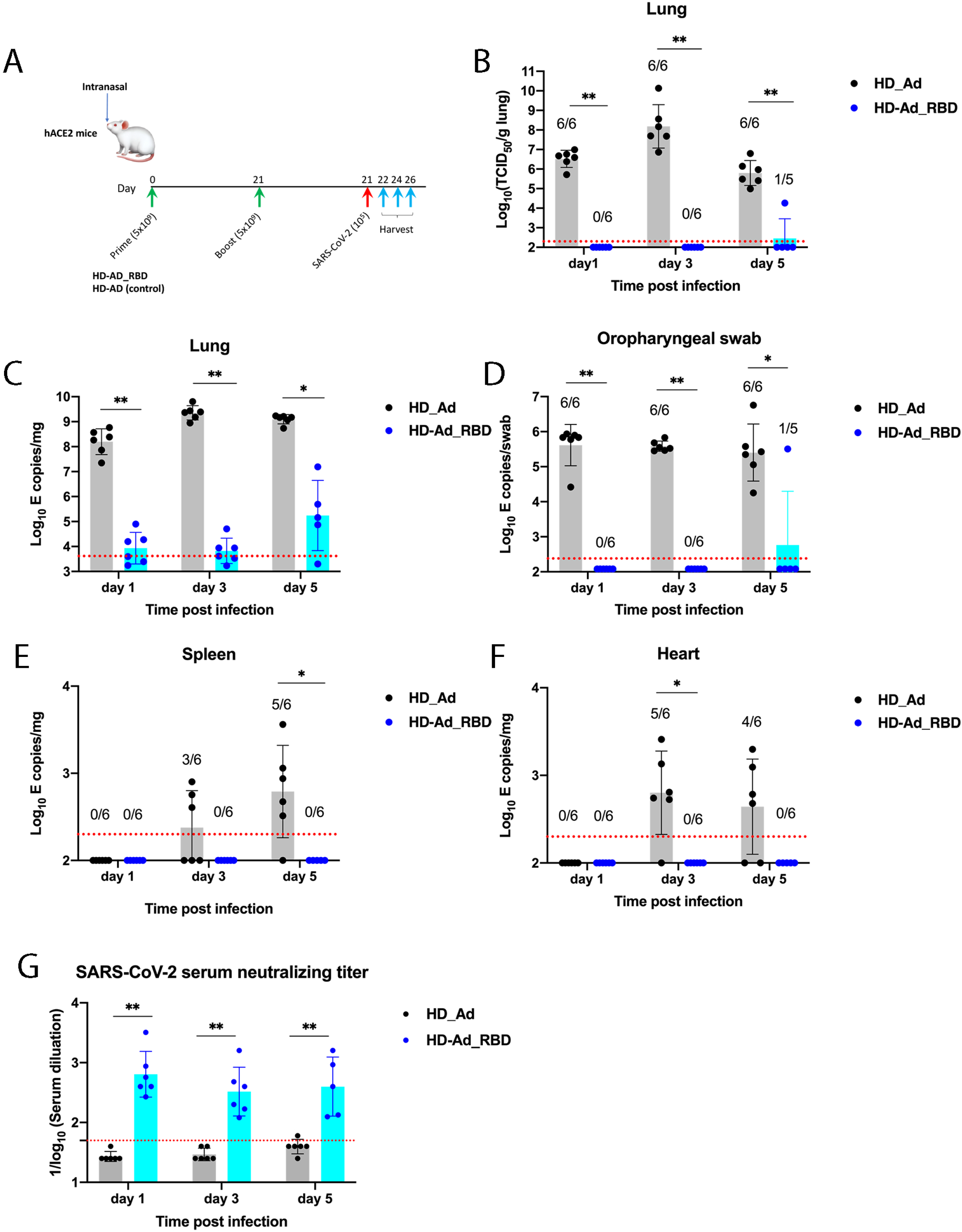
HD-Ad_RBD protects the upper and lower respiratory tracts against SARS-CoV-2 infection. (**A**) **Scheme of the experiment**. hACE2 mice were immunized intranasally with HD-Ad_RBD by prime-boost regimen at a dose of 5×10^9^ viral particles. Three weeks after the second vaccination, the animals were challenged intranasally with 10^5^ TCID_50_ of SARS-CoV-2. At day 1, 3 and 5 post-infection, mice were euthanized and lungs, spleen and heart were harvested for viral burden and cytokine analysis. (**B**) **Titers of infectious SARS-CoV-2**. The number of infectious virus in lungs were determined by CPE assays. (**C-F**) **RNA levels of SARS-CoV-2 determined by qRT-PCR**. Viral RNA levels in the lung, spleen, heart, and oropharyngeal swabs were measured at indicated time points by qRT-PCR. (**G**) **Neutralizing activity of sera against SARS-CoV-2.** Sera from mice at different time points after SARS-CoV-2 infection were collected and the levels of neutralizing antibody were measured. Each dot represents an animal. Bars and errors represent the geometric mean with geometric SD. The red dotted lines indicate the limit of detection (LOD) of assays. Statistical analyses were performed by Mann-Whitney test: *p < 0.05; **p <0.01; ***p < 0.001.

Notably, there were no detectable infectious virus in the lungs of 16 mice immunized with HD-Ad_RBD as determined by the TCID_50_ assay, whereas high levels of the infectious virus were detected in all mice (*n*=18) vaccinated with the vector control (**Fig. 3B**). Only one mouse immunized with HD-Ad_RBD showed a detectable but low level of infectious virus, which may be an outlier due to improper vaccination. We noticed that mice sometimes sneezed out the vaccine solution during intranasal inoculation. Using the primers that are specific to the sequence of the SARS-CoV-2 *E* gene, we detected very high levels of viral RNA (10^8^ to 10^9^ copies/mg) in the lungs of mice vaccinated with the HD-Ad vector control. In contrast, the viral RNA levels were reduced by >4 log_10_ in 16 out of 17 mice vaccinated with HD-Ad_RBD (**Fig. 3C**). The very low RNA levels in the lungs of mice vaccinated with HD-AD_RBD may reflect the input and nonreplicating virus. This is consistent with two observations. First, comparing mice of the control group at day 1 and 3 post-infection, there was a significant increase (>1 log_10_) in both the infectious viral titer and the viral RNA copy number in the lungs at day 3 compared to day 1, indicating substantial viral replication in these mice (**Fig. 3B & C**). However, this trend was not observed in the mice vaccinated with HD-Ad_RBD as no infectious virus was detected in these mice and the viral RNA level remained constant at day 1 and 3 post-infection (**Fig. 3B & C**). Second, similar low SARS-CoV-2 RNA levels (~10^4^ copies/mg) were observed at these time points in BALB/c and C57BL/6 mice lacking hACE2 receptor expression, which represent the input, nonreplicating virus^37^.

Remarkably, intranasal delivery of HD-Ad_RBD provided effective protection of the upper respiratory tract as judged by the absence of measurable viral RNA in the oropharyngeal swabs (**Fig. 3D**). Only one mouse vaccinated with HD-Ad_RBD showed a detectable level of viral RNA, which is the same abovementioned mouse that showed infection in the lungs. All mice in the control group exhibited high levels of viral RNA (10^6^ copies/swab). We also detected no measurable or very low levels of viral RNA in the heart and spleen of mice vaccinated with HD-Ad_RBD (**Fig. 3E & F**).

The protection of HD-Ad_RBD vaccinated animals against SARS-CoV-2 is likely due to high neutralizing antibody levels. To test this, we collected the sera of these viral challenged animals and measured the neutralizing antibody levels. As expected, since the sera were collected shortly after the challenge by SARS-CoV-2 (day 1 to 5 post-infection), the control group mice did not have enough time to develop detectable levels of neutralizing antibody (**Fig. 3G**). In animals vaccinated with HD-Ad_RBD, high levels of neutralizing antibody against SARS-CoV-2 were detected, with the reciprocal ID_50_ GMTs ranging from 328-640 (**Fig. 3G**), which is in the same range of neutralizing antibody detected in the vaccinated but not challenged animals (**Fig. 2G**).

### HD-Ad_RBD vaccination protects mice against SARS-CoV-2-induced inflammation in the lung

We next examined the effect of the vaccine on lung inflammation. The lung tissues at day 3 post-infection were chosen for this analysis and the mRNA levels of several proinflammatory cytokines and chemokines were measured and normalized against the same samples prepared from the naïve mice, which were mice that were vaccinated with the HD-Ad vector control but not challenged by SARS-CoV-2.

In the mice that were vaccinated with the HD-Ad vector control and challenged by SARS-CoV-2, there was a dramatic increase on the mRNA levels of *IL-6*, *CXCL10*, and *CXCL11* (**Fig. 4**). Remarkably, in mice that were vaccinated with the HD-Ad_RBD and challenged by SARS-CoV-2, these proinflammatory cytokines and chemokines were reduced to the levels of naïve mice. The mRNA levels of *CXCL1*, *IFN-γ*, *IL-1β* and *IL-11* were also significantly lower in the lung tissues of animals immunized with HD-Ad_RBD compared to the HD-AD control (**Fig. 4**).

**Figure 4.**
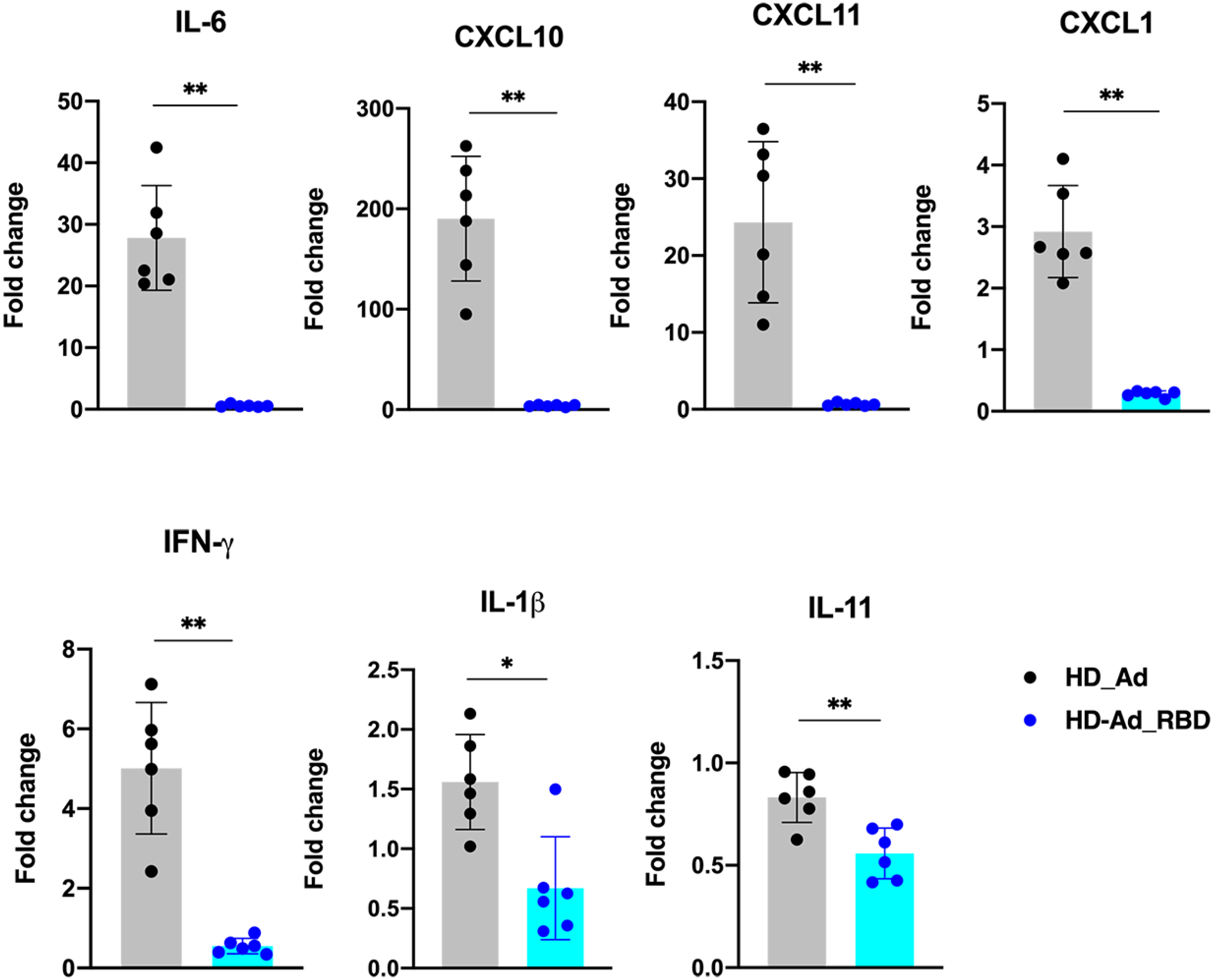
HD-AD_RBD prevents inflammation in the lung from viral challenge. Fold change in gene expression of indicated cytokines and chemokines from lung homogenates at 3-day post infection was determined by qRT-PCR after normalization to *GAPDH* levels and comparison with the naive unchallenged control. Each dot represents an animal. Bars and errors represent the mean with SD. Statistical analyses were performed by Mann-Whitney test: *p < 0.05; **p <0.01; ***p < 0.001.

Taken together, our results demonstrate that immunization with HD-Ad_RBD decreases both viral infection and consequent inflammation in the lungs of animals infected with SARS-CoV-2.

## Discussion

Although several COVID-19 vaccines have been approved for emergency use in humans, they have typically not provided effective protection against viral infection in the upper airway of animals or humans. The ChAdOx1 nCoV-19 vaccine, developed by the University of Oxford, and AstraZeneca, is based on a replication-incompetent chimpanzee adenovirus^15^. This vaccine was tested in rhesus macaques in a prime-boost regimen (4 weeks apart), at a dose of 2.5×10^10^ viral particles administered intramuscularly. The animals developed moderate neutralizing antibody titers (reciprocal ID_50_ GMT:10-160, after the boost) and were mostly protected from lung disease. However, the viral replication in the upper respiratory tract was not controlled^15^. The mRNA-1273 vaccine developed by the National Institutes of Health (NIH) and Moderna was tested at two doses (10 μg and 100 μg) in rhesus macaques in a prime-boost regimen with a 4 week-interval by intramuscular injection^7^. For the high dose case, the vaccine induced high levels of IgG and neutralizing antibody, reaching a reciprocal GMT of 36,386 and a reciprocal ID_50_ GMT of 1,862, respectively, after the boost vaccination. Moreover, the animals were almost completely protected from SARS-CoV-2 infection in the lower respiratory tract (BALs). However, even in the high-dose group, viral RNA was detected in the upper respiratory tract (nasal swabs) in three out of eight macaques on day 1 post-infection and one out of eight on day 4 post-infection^7^. A similar finding was observed when the mRNA-1273 vaccine was tested in mice^38^. The BNT162b2 mRNA vaccine developed by BioNTech and Pfizer was also tested in rhesus macaques at two doses (30 μg and 100 μg) in a prime-boost regimen by intramuscular injection^39^. High levels of neutralizing antibody (reciprocal ID_50_ GMT: 1,689) were detected in the high-dose group after boost vaccination. These animals were completely protected against SARS-CoV-2 infection in BALs. However, viral RNA was detected in high levels in nasal swabs of five out of six vaccinated animals at day 1 post-infection, and in oropharyngeal swabs of three out of six vaccinated animals at day 1 post-infection, and in two out of six vaccinated animals at day 3 post-infection^39^.

Our HD-Ad_RBD vaccine confers a complete protection against SARS-CoV-2 infection in the upper air way of the hACE2 mice, as evidenced by the absence of viral RNA almost immediately after the intranasal infection with SARS-CoV-2 (day 1 post-infection). We also did not detect any infectious virus in the lungs at day 1 post-infection. These results show a rapid control of viral replication almost immediately after viral infection. Taken together, our data suggest that HD-Ad_RBD has the potential to control infection at the site of inoculation, which should not only prevent virus-induced disease but also transmission.

The superior protection efficacy of our HD-Ad_RBD vaccine is likely due to high antigen expression levels and the intranasal delivery route employed. Replication-deficient adenoviral vectors with one or more early genes deleted (the first- and second-generation Ad vectors) are among the most efficient vehicles for *in vivo* gene delivery and they have been used for the development of COVID-19 vaccines. However, one of the major limitations in the use of these vectors for therapeutic purposes is the transient nature of the transgene expression and the systemic toxicity, which are both due to inflammatory and immune responses triggered by the residual expression of viral proteins^40–42^. To overcome these problems, HD-Ad, the third generation of Ad vectors, with all viral coding sequences deleted, was used in our study. HD-Ad minimizes the host immune response to the vector and allows long-term expression of the transgene in host tissues or organs, thereby providing a safer and more efficient way for gene delivery and transgene expression^30–34^. Consistently, we detected high level expression of the RBD in transfected cells and the majority of the protein was secreted, which ensures efficient antigen-dependent stimulation of the host immune system. Consequently, the HD-Ad_RBD vaccine induced extremely high levels of antigen-specific antibody response. For example, the IgG reciprocal GMT in sera reached 1,837,920 even though we used a moderate dose of the vaccine (5×10^9^ viral particles) for vaccination. We also detected high levels of neutralizing antibody against the SARS-CoV-2 virus, and the reciprocal ID_50_ GMT was close to 1,000, which is in the same range of the neutralizing antibody induced by the two mRNA vaccines (mRNA-1273, BNT162b2) described above.

The superior safety profile of HD-Ad allows us to deliver the vaccine by an intranasal route. Compared to intramuscular injection, immunization through the nasopharyngeal route has the advantage of eliciting a local immune response, including secretory IgA antibodies that confer protection at or near the site of infection of respiratory pathogens^43^. Consistently, high levels of antigen-specific IgA were detected in both sera and BALs of the animals vaccinated with HD-Ad_RBD, indicating that this vaccine induced robust mucosal immunity, in addition to strong systemic immunity.

To our knowledge, there has been only one published study evaluating the intranasal delivery of a COVID-19 vaccine candidate. Hassan *et al.* compared intramuscular injection and intranasal delivery of a chimpanzee Ad-based (simian Ad-36) SARS-CoV-2 vaccine in mice^12^. They found that intranasal delivery, but not intramuscular injection, of the vaccine induced the production of IgA in BALs and the sera of the animals. In K18-hACE2 mice, a single intranasal delivery of the vaccine (10^10^ viral particles) followed by challenge with SARS-CoV-2 resulted in very low levels of viral RNA in the nasal turbinates or washes, suggesting almost complete protection of the upper respiratory tract against infection^12^. One caveat of this experiment is that they used a lower dose of SARS-CoV-2 (10^3^ foci forming units) to challenge the single intranasal vaccinated hACE2 mice, as compared to the 4×10^5^ FFUs used for other experiments in their studies^12^.

Key to controlling COVID-19 will be the development of vaccines that provide long-term protection, a property requiring long-term T cell responses^44^. It is known that adenovirus-based vaccines can elicit strong T cell memory^45^ and since our HD-Ad vaccine shares the same capsid proteins as regular adenovirus-based vaccines, we expect long-term T cell memory from our HD-Ad_RBD vaccine as well. Indeed, spleen cell samples from mice 6.5 months after a single-dose (5×10^9^ viral particles) vaccination with HD-Ad_RBD show high levels of antigen-specific IFN-γ (~14,000 pg/ml), an indication that our vaccine elicits a strong and long-term T cell response.

In additional to its superior safety profile, HD-Ad also has a large cloning capacity (up to 36 kb) for transgenes and this makes it possible to deliver large genes or multiple genes. These features, together with the remarkable success of the HD-Ad_RBD vaccine demonstrated in this study, make HD-Ad an ideal platform for the construction of multivalent vaccines targeting SARS-CoV-2 and its emerging variants, work which is now underway in our laboratories.

## Materials and Methods

### Virus and animals

The SARS-CoV-2 virus was isolated from local patients at Toronto in March 2020. All work with infectious SARS-CoV-2 was performed in the Containment Level 3 (CL-3) facilities at University of Toronto using appropriate protective equipment and procedures approved by the Institutional Biosafety Committee.

BALB/c and K18-hACE2 C57BL/6 mice were purchased from The Jackson Laboratory. K18-hACE2 mice were bred in house and each mouse was genotyped before use.

### Ethics Statement

All of the animal procedures were approved by the University of Toronto Animal Care Committee (Animal Use Protocol# 20012653) and Hospital for Sick Children (Animal Use Protocol #1000057758). All experimental procedures were performed in accordance with the Canadian Council on Animal Care (CCAC) and University of Toronto and Hospital for Sick Children regulations.

### Construction of HD-Ad_RBD

A viral vector genome expressing a secreted form of the RBD of SARS-CoV-2 spike protein was constructed by multiple steps. We started with a plasmid, which is based on pBluscript containing the chicken beta actin gene (CBA) promoter and the ploy A signal from bovine growth gene (BGHpA), by inserting UbC gene intron 1 into the plasmid between CBA promoter and BGHpA by In-fusion cloning. To build a plasmid expressing RBD from the CBA promoter, we inserted the RBD sequence with the signal peptide of the human cystatin S gene at its 5’ end after the UbC intron between the restriction sites, EcoRV and ApaI. Finally, we inserted the RBD-expression cassette as AscI fragment from the RBD expressing plasmid into viral vector, pC4HSU-NarD at AscI site, resulting in the new plasmid, pC4HSU-NarD-RBD for the vaccine production.

### Vaccine production

The vaccine production was carried out using 116 cells^46^. A helper-virus, NG163, was used to provide vector DNA replication and the production of viral capsid proteins. The packaging signal sequence in the helper-virus was flanked by two loxP sites. During vector production, the host cells expressed the Cre recombinase which cleaved off the packaging signal of the helper virus. Thus, only HD-Ad vector particles were assembled. The large-scale production of HD-Ad vectors was carried out in suspension cells using 3L Bioreactors. The vector particles were harvested from the cell lysate and purified through two rounds of CsCl gradient centrifugation.

### Immunization of BALB/c mice

Female BALB/c mice were intranasally immunized with different doses (10^8^ to 10^10^ viral particles) of HD-Ad_RBD or the HD-Ad vector control (HD-C4HSU) in 20 μl of PBS containing 40 μg/ml of DEAE-Dextran and 0.1% LPC. Depending on the immunization regimen, the mice were either euthanized or boosted with the vaccine after 3 weeks. The boosted mice were sacrificed at 3 weeks after the second vaccination. Mouse tissues as well as samples of blood and bronchoalveolar lavage fluids (BALF) were collected.

### ELISA

Purified RBD protein was used to coat the flat-bottom 96-well plates (Thermo Scientific NUNC-MaxiSorp) at a concentration of 1 μg/ml in 50 mM carbonate coating buffer (pH 9.6) at 4 °C overnight. The following day, plates were blocked with solution containing 1% BSA in PBST. Serially diluted mouse sera or BALF were added and incubated at 37 °C for 1 h. Antibodies including goat anti-mouse IgG horseradish peroxidase (HRP)-conjugated (31430 thermo) and anti-mouse IgA HRP-conjugated (626720 Invitrogen) were diluted 1:5000, or 1:200 in blocking solution. After incubation for 1 h, the plates were washed and developed with 3,3’,5,5’-tetramethylbiphenyldiamine (TMB, 34028, Thermo) for 10 min. The reactions were stopped with 1.0 M H_2_SO_4_ stop solution. The absorbance was measured on a microplate reader at 450 nm (A450). The endpoints of serum or BALF dilutions were calculated with curve fit analysis of optical density values for serially diluted serum or BALF with a cut-off value set to two to three times the background signal.

### Flow cytometry analysis

Left lungs from HD-Ad_RBD vaccinated and control mice were harvested and digested for 45 min at 37 °C in digestion buffer containing liberase 2 μg/ml and Type IV DNase I 25 units/ml. The lung infiltrated cells were cultured with purified RBD protein at 10 μg/ml for 12 h at 37 °C followed by a 6 h treatment with GolgiPlug (BD 555028). After blocking with FcγIII and FcγII receptors antibody (BD Pharmingen, 553142), cells were stained with live/dead fixable cell stain (Invitrogen 34955), CD44 BV510, CD4 BV711, and CD8a APC-Cy™7 (BD) antibodies. Stained cells were fixed and permeabilized with Cytofix/Cytoperm Fixation/Permeabilization (BD 555028), and then intracellularly stained with anti-IFN-γ APC (BD). Cells were analysed on a Becton Dickinson LSR II CFI (SickKids Flow Cytometry Facility), using Flowjo x 10.0 software.

### SARS-CoV-2 neutralization assay

Heat-inactivated serum was serial diluted (2-fold) in DMEM and incubated with 200 TCID_50_ of SARS-CoV-2 for 2 h at 37°C. For each dilution, there were six technical replicates. After the 2h neutralizing, the serum-virus mixture was then incubated with 20,000 Vero-E6 cells supplied with 2% FBS at 37°C. CPE of each well was examined at day 5. The highest dilution of serum that can protect 50% of cells from SARS-CoV-2 infection is considered as the neutralizing antibody titer, as described in^4^.

### SARS-CoV-2 infection of hACE2 mice

Six- to eight-week-old K18-hACE2 C57BL/6 mice (female and male at equal ratio) were intranasally immunized with 5 × 10^9^ viral particles of HD-Ad_RBD or the HD-Ad vector control (HD-C4HSU) in 20 μl of PBS containing 40 μg/ml of DEAE-Dextran and 0.1% LPC. The animals were boosted with the same dosage of HD-Ad_RBD or HD-Ad, respectively, three weeks after the prime vaccination. Three weeks after the boosted vaccination, mice were infected with 1×10^5^ TCID_50_ of SARS-CoV-2 in 50 μl of DMEM via the intranasal route. Animal were euthanized at day 1, 3 and 5 post-challenge and samples were collected for further analysis.

### Measurement of viral burden

The infectious virus number was determined by a cytopathogenic efficiency (CPE) assay. Vero-E6 cells (30,000) were seeded into a 96-well plate one day before the inoculation. Collected tissues were weighed and homogenized with stainless steel beads (Qiagen, #69989) in 1 ml of DMEM with 2% FBS. Lung homogenates were centrifugated at 3,000 g for 5 min and the supernatants were collected. Serial 10-fold dilutions of the lung homogenates were then added into the monolayer Vero-E6 cells. For each dilution, there were six technical replicates. After 5 days of culture, the CPE of each well was examined, and the virus titer (TCID_50_) was calculated according to the Karber method^47^ and normalized by the organ weight.

The viral RNA copy number was measured by one-step real-time quantitative PCR (qRT-PCR) as described in^48^. Briefly, collected tissues were weighted and homogenized with stainless steel beads (Qiagen, #69989) in 1ml of Buffer RLT. The RNA was extracted with the RNeasy Mini kit (Qiagen, #74104). For oropharyngeal samples, swabs were eluted in 500 μl PBS by vortex, and the viral RNA in the PBS eluent was extracted using QlAamp Viral Mini kit (Qiagen, #52906). Primers and TaqMan probes (IDT, #10006890, 10006891 and 10006893) that target the SARS-CoV-2 envelop (*E*) gene were used to detect the genomic/subgenomic viral RNA. The standard curve of Cq-value to viral copy number was generated using serial 10-fold dilutions of the *E* gene plasmid DNA template (IDT, #10006896). qRT-PCR was performed with the NEB Luna universal probe kit (E3006) under the following reaction conditions: 55°C for 10 min, 95°C for 1 min, and 40 cycles of 95 °C for 10 sec and 58 °C for 30 sec. The viral RNA copies were determined by converting the Cq-value according to the standard curve.

### Measurement of cytokine and chemokine mRNA levels

RNA of the lung homogenates was extracted and qRT-PCR were performed as mentioned above. Primers used in this experiment were listed in Table S2. The mRNA level of cytokines and chemokines were normalized to *GAPDH*. Fold change was calculated using the 2^−ΔΔCq^ method by comparing SARS-CoV-2 infected mice to uninfected mice.

## Acknowledgement

This work was supported by Canadian Institutes of Health Research (CIHR) grants VR1-172771 and VS-1-17553138 (to JL, JH and JR).

---

**Table S1.**
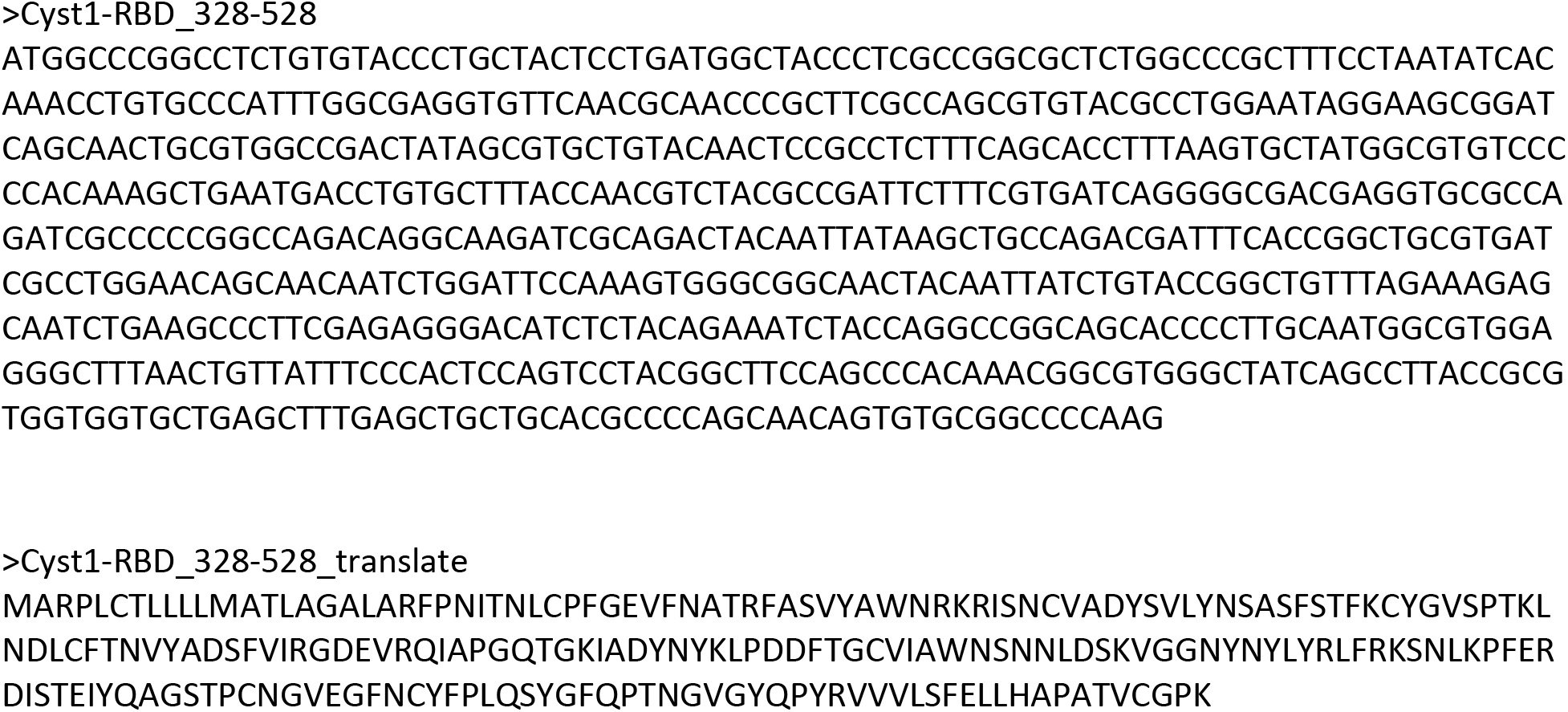
Codon optimized RBD (328-528) of SARS-CoV-2 Spike Protein.

**Table S2.**
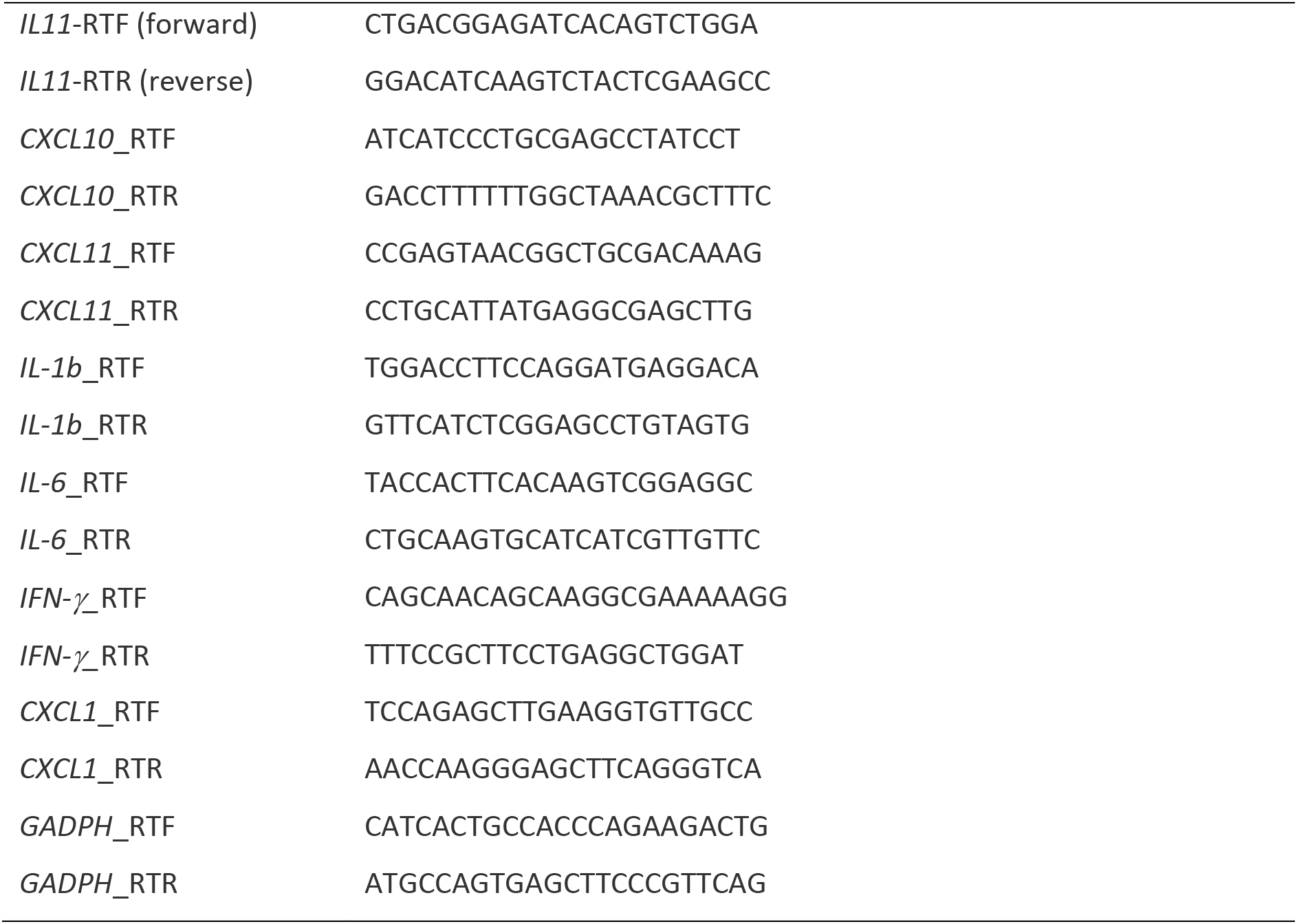
Primers used in qRT-PCR analysis of cytokine and cytokine and chemokine mRNAs.

